# Effects of incrementally increased plant-based protein intake on gut microbiota and inflammatory–metabolic biomarkers in healthy adults

**DOI:** 10.1101/2025.10.09.680873

**Authors:** Samira B. R. Prado, Annalena Kamm, Katharina Dannenberg, Isabel Keidel, Victor C. Castro-Alves, Tuulia Hyötyläinen, Marleen Lentjes, Dirk Repsilber, Tatiana M. Marques, Robert J. Brummer

## Abstract

Shifting to a plant-based diet naturally alters protein source choices. In many countries, protein from yellow pea is widely used as main ingredient in meat alternatives. Still, its biological effects, especially regarding gastrointestinal health, remain incompletely understood. The aim of our study was to investigate how a weekly increase in the intake of a well-characterized pea protein isolate affects surrogate markers of health, fecal short-chain fatty acids and gut microbiota composition in healthy individuals. Male and female adults (N=29) participated in this exploratory intervention study. A 4-week pre-intervention period for questionnaires and fecal samples collection was followed by a 4-week supplementation. Participants consumed isolated pea protein in weekly increasing amounts, starting from 0.25 g/kg body mass/day in week 5 to 1.00 g/kg body mass/day in week 8. Questionnaire data, fecal samples as well as fasting blood and 24-h urine samples were collected weekly. Data from biological samples and questionnaires confirmed a healthy study population and compliance. Fecal calprotectin levels significantly increased only in a subset of participants, which was also accompanied by higher fecal water cytotoxicity *in vitro*. Short-chain fatty acids mainly rose in those subjects with stable calprotectin levels. Relative abundances of *Limosilactobacillus frumenti, Odoribacter splanchnicus* and *Lactobacillus crispatus* increased significantly in the total population during the intervention while the relative abundance of *Bifidobacterium longum* and *Bifidobacterium catenulatum* decreased. Our results indicate that an increased intake of pea protein isolate affects the growth of certain beneficial bacteria strains and differentially influences markers related to gut inflammation in healthy individuals.

## 1. Introduction

A societal shift from animal to plant-based diet is taking place, driven in part by environmental concerns (Abe-Inge et al., 2024). Pulses – the edible seeds of legumes, including beans, peas, and lentils – as a plant-based protein have been shown to support cardiovascular, metabolic, and anti-inflammatory health (Luzardo-Ocampo & Gonzalez de Mejia, 2025). Among these, peas have emerged as a prominent source due to their adaptability to grow in moderate climates and their lower allergenic potential compared to soy (Benković et al., 2023). Additionally, they have good emulsifying properties, the ability to form gels (Broucke et al., 2025), and a desirable nutritional quality regarding amino acid profile (Auer et al., 2024). These attributes may explain the growing investigation of isolated pea protein in Europe, as well as its application in various food products (Czaja et al., 2025; Ebert et al., 2021; Moll et al., 2023) – for example, commercially available meat alternatives (Plattner et al., 2024; Zhu et al., 2021). Plant-based proteins often have a lower digestibility compared to animal proteins, wherefore they may escape the digestion and absorption in the small intestine, especially when consumed in higher amounts, reaching the colon and undergoing microbial colonic fermentation (Boven et al., 2024). Proteolytic fermentation in the gut is often associated with detrimental effects (Windey et al., 2012a), for example, due to the production of ammonia (Levitt & Levitt, 2018), which has been associated with intestinal inflammation, intestinal barrier dysfunction, and increased risk of colorectal cancer (Blachier et al., 2007; Fung et al., 2013; Windey et al., 2012a). In contrast, short-chain fatty acids (SCFA), which are also a product of colonic fermentation, are generally considered beneficial for gut health by supporting lipid, glucose, and immune homeostasis (Campos-Perez & Martinez-Lopez, 2021; Yao et al., 2022). Among SCFA, acetate, propionate, and butyrate are primarily produced from dietary fiber fermentation (Yao et al., 2022), whereas higher protein intake has been related with greater levels of propionate and valerate (Mak et al., 2025).

Drastic changes in dietary protein sources can furthermore affect gut microbiota composition and function, potentially impacting human health (Alvarenga et al., 2024; David et al., 2014; Wu et al., 2022). In a systematic review and network meta-analysis, quantity and source of protein were not related with changes in microbiota composition, but higher dietary protein intake increased microbiota-derived metabolites linked with proteolytic fermentation (Mak et al., 2025). So far, only *in vitro* fermentation studies have been performed using isolated pea protein (Karlsson et al., 2024) and data on *in vivo* effects in humans are scarce.

Non-direct microbiota related compounds are also involved in gut inflammation. For instance, fecal calprotectin, which is produced by neutrophils and a surrogate marker for intestinal inflammation (Jukic et al., 2021; Stríz & Trebichavský, 2004). Animal-based diets have been associated with its increased levels in patients with Crohn’s disease and ulcerative colitis (Bolte et al., 2021). In patients with type 2 diabetes, only plant-based and not animal-based protein increased calprotectin in blood (Markova et al., 2020). In healthy individuals it is unclear whether high plant-based protein intake would affect such markers.

To the best of our knowledge, no human dietary intervention study has yet investigated the association between dietary isolate pea protein intake and gut-related markers of health in a non-diseased population. Therefore, the aim of this single-arm exploratory dietary intervention study was to evaluate the effect of an increasing intake of isolated pea protein on various surrogate markers of health, fecal SCFA, and gut microbiota composition in healthy individuals.

## 2. Materials and Methods

### 2.1. Study Design

This intervention study was conducted at Örebro University, Sweden in two periods between October–December 2021 and February–April 2022, respectively. The study was approved by the Swedish Ethical Review Authority (Dnr 2021-03256, 08-24-2021), conducted in accordance with the Declaration of Helsinki, and registered at www.clinicaltrials.gov (NCT05367804).

After assessing eligibility on a first screening visit and after participants have signed an informed consent, participants came to the study center on eight occasions scheduled one week apart (**Figure 1**). On the first three study visits, they handed in fecal samples collected at home or at the study site, a completed Gastrointestinal Symptom Rating Scale (GSRS), an International Physical Activity Questionnaire (IPAQ), a Bristol Stool Scale (BSS) diary as well as three 24-h food diaries. Additionally, participants were weighted on visit 3 to calculate the amount of the protein supplement they would receive from visit 4 onwards. Study visits 4–8 were scheduled in the morning after an overnight fast of at least 10 hours and included, in addition to fecal sample collection and completion of questionnaires, the collection of 24-h urine samples as well as fasting blood samples.

**Figure 1.**
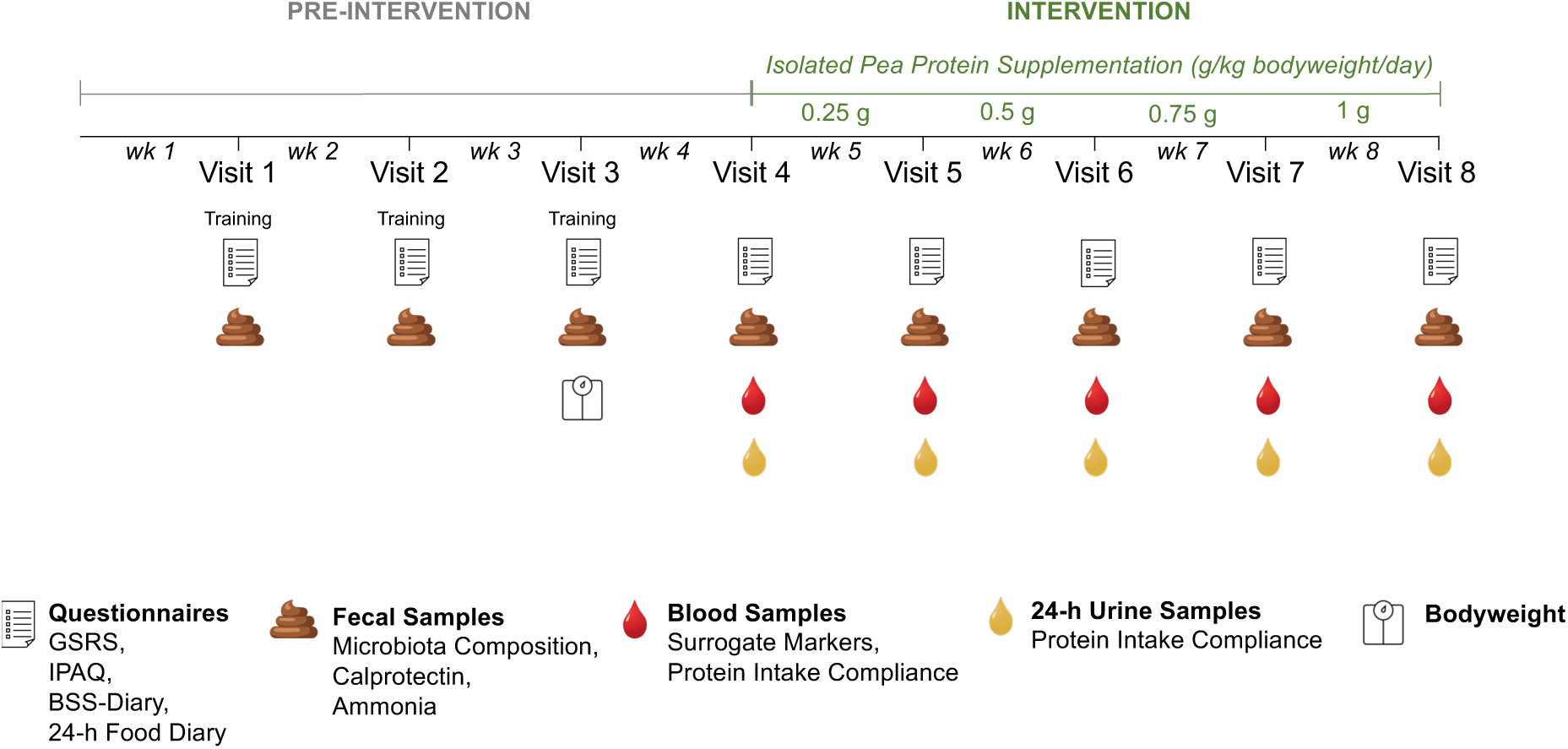
Study Design Summary. GSRS: Gastrointestinal Symptom Rating Scale, IPAQ: International Physical Activity Questionnaire, BSS: Bristol Stool Scale. Questionnaires in the pre-intervention period were filled in as training for the intervention period.

### 2.2. Study Subjects

Healthy male and female adults were recruited via social media, the university’s webpage as well as posters placed in public areas in Örebro and the University Campus. Applied inclusion criteria were: age 18–45 years; body mass index (BMI) between 18.5–30 kg/m^2^; stable weight within the previous three months; maintenance of the usual physical activity habits during the study; intake of dietary fiber between 15–25 g/day (as evaluated by two food diaries and one 24-h recall); omnivores or vegetarians.

Interested individuals were excluded if they fulfilled any of the following exclusion criteria: acute or chronic disease, inflammatory or functional gastrointestinal diseases and any other disease or disorder that could affect the outcome of the study; use of a medication that may interfere the study outcome; eating disorder; high-protein intake (more than 15 % of energy or maximum 1.2 g/kg body mass/day as evaluated by two food diaries and one 24-h recall); use of antibiotic medication during the last three months prior the first visit; use of laxative or anti-diarrheal medication within the past three months before the study; regular consumption of probiotic or prebiotic products for the past six weeks before the study; special diet that is considered to affect the study participation and/or study results, for example, high-protein diets; more than five hours of moderate-vigorous exercise/week; pregnancy or breastfeeding; intolerance to dietary supplements that will be used in the study; tobacco and nicotine use; abuse of alcohol or drugs.

### 2.3. Study Product

During the intervention period (visit 4–8), participants consumed protein isolate from yellow pea (Pisane C9, COSUCRA, Belgium), which has been well characterized in the literature (Auer et al., 2024; Broucke et al., 2025; Schumacher et al., 2025; Tiong et al., 2024), and is widely used as food product ingredient (Czaja et al., 2025; Ebert et al., 2021; Moll et al., 2023). The protein isolate was manufactured by dehulling and milling the peas, solubilization in water, and decantation to separate the soluble (protein) and insoluble (fiber, starch) fractions, pasteurization of the soluble protein fraction which was followed by purification, concentrating, and spray-drying (Moll et al., 2023). An isolate was used in the study to minimize the impact of antinutritional factors on the gastrointestinal system and minimize confounders with other food components from the intervention product (Ashkar & Wu, 2023; Auer et al., 2024). The chemical composition of the provided isolated pea protein was calculated based on a study within our research consortium ((Auer et al., 2024); **Supplementary Table 1**).

The amount of protein powder was calculated based on the individual body mass at study visit 3 and increased gradually each week (0.25, 0.50, 0.75 and 1.00 g/kg body mass/day). The protein content of the supplement was provided by the product specification (83.6 g/100 g of powder). The protein powder was weighed, and the daily amount was packaged in three equal portions which participants were asked to ingest together with breakfast, lunch, and dinner to distribute the protein intake over the day.

### 2.4. Biological Samples

Plasma was used to analyze C-reactive protein (CRP), triglycerides, creatinine, urea, as well as total, high-density lipoprotein (HDL) and low-density lipoprotein (LDL) cholesterol, while serum was analyzed for insulin. Whole blood was used to measure glucose levels.

Urine samples were collected in a designated container (Sarstedt, Germany) by the participants during 24 h. The collection period started in the morning on the day before the study visit, excluding the morning urine, and lasted until the morning on the day of the study visit including the morning urine. Between the urine sampling occasions, participants were instructed to store the container at 6–8 °C. Missing sampling occasions resulted in exclusion of the data. Aliquots from the homogenized container were collected with vacuum tubes (Sarstedt, Germany), immediately stored at -20 °C, and further transferred to -80 °C until analysis for creatinine, urea, and uric acid.

Fecal samples were collected by the participants up to one day before the study visit using an EasySampler Stool Collector Kit (GP Medical Devices, Denmark) and fecal sample collection tubes (Sarstedt, Germany). Participants were instructed to store the tubes at -20 °C and samples were transported to the study center in a frozen transport container (Sarstedt, Germany). The tubes were immediately stored at -20 °C and further transferred to -80 °C until analysis. For fecal calprotectin analysis, samples were thawed at room temperature and extracted using CALEX cap tubes (BÜHLMANN, Switzerland).

All biological samples mentioned above were analyzed at the Laboratory Medicine Clinic (Clinical Chemistry Department, Örebro University Hospital) using clinically automated procedures.

Fecal ammonia concentrations were determined using the commercially available Ammonia Kit (Megazyme, Ireland) according to the manufacturer’s instructions.

Fecal samples were also used for measuring relative levels of the SCFA butyrate, acetate, propionate, and valerate by ultra-high performance liquid chromatography coupled to time-of-flight high-resolution mass spectrometry (UHPLC-Q-TOF-HRMS) after derivatization of fecal extracts using 3-nitrophenylhydrazine (3-NPH) (Seeburger et al., 2023). Briefly, 20 mg of fecal sample was mixed with 100 µL cold methanol containing internal standards (10 µg/mL each): aceticacid-d4, butyric acid-d8 and propionic acid-d2. After ultrasonication (5 min), the extract was centrifuged (10000 *g*, 5 min, 4 °C) and a 50 µL-aliquot transferred to a vial. Each sample was further derivatized for 1 h using 50 mM 3-NPH, 50 mM N-ethylcarbodiimide and 7 % (v/v) pyridine in aqueous methanol solution. After incubation, 0.2 % formic acid was added to stop the derivatization reaction, and the sample was immediately analyzed using a 1290 UHPLC-Q-TOF system (Agilent, USA) equipped using an Acquity BEH C18 column (2.1 × 100 mm, 1.7 µm; Waters Corporation, USA). Mobile phase (A) 0.1 % formic acid in water and (B) acetonitrile were eluted at 0.4 mL/min. The injection volume was 5 µL and autosampler and column temperature were 10 °C and 50 °C, respectively. Data was acquired in negative ion mode. MassHunter Workstation Software (Agilent, USA) was used for data acquisition and processing.

### 2.5. Microbiota Analysis

The extraction of DNA and NGS sequencing for the fecal microbiota analysis was performed at Clinical Genomics, Örebro using the QIAsymphony PowerFecal Pro DNA Kit (Qiagen, Germany) on a QIAsymphony SP liquid handler (Qiagen, Germany) following the manufacturer’s instructions, with a few minor modifications to the pretreatment step. Briefly, aliquots of approximately the size of a pea were collected from each fecal sample using a sterile 10 µL plastic loop and placed into PowerBead Pro Tubes (Qiagen, Germany) containing 750 µL CD1. Bead beating was performed on a FastPrep 24 bead beater (MP Biomedicals, USA) for 1 min at 6 m/s. After centrifugation (15000 *g*, 1 min), the supernatant was transferred to new 2 mL microcentrifuge tubes. Proteinase K digestion was performed by incubation with 30 µL Proteinase K (20 mg/mL) for 30 min at 56 °C. After digestion, 300 µL CD2 was added, samples were centrifuged (15000 *g*, 1 min) and the supernatant was transferred to new 2 mL micro tubes which were loaded on the QIAsymphony. The elution volume was 100 µL. The purified DNA was quantified on a Qubit 2.0 fluorometer (Thermo Fisher, USA) using the Qubit 1X dsDNA HS Assay Kit (Thermo Fisher, USA).

Library preparation was performed on NGS STAR (Hamilton, USA) using the Illumina DNA Prep kit and Illumina DNA/RNA UD Indexes (Illumina, USA) according to the manufacturer’s instructions. The libraries were quantified on a Qubit 3.0 fluorometer (Thermo Fisher, USA) using the Qubit 1X dsDNA HS Assay Kit (Thermo Fisher, USA) and average fragment lengths determined using the 4200 Tapestation system (Agilent, USA) with the High Sensitivity D5000 screentapes and reagents (Agilent, USA). Sequencing was performed on a NextSeq 2000 sequencer using NextSeq 2000 P3 Reagents (300 Cycles) (Illumina, USA) and sequences from two runs were combined for each sample to gain sufficient read depth. Yielded data was subsequently processed with default settings of the taxprofiler pipeline (version 1.1.2; nextflow version 23.10.0) (Stamouli et al., 2023) using the MetaPhlAn (4.0.6) taxonomic profiler.

### 2.6. Questionnaires

During the screening visit, participants were interviewed to verify if they met the study’s inclusion and exclusion criteria. Additionally, a 24-h dietary recall was performed together with the participant by trained staff. Participants were also instructed and trained on how to properly fill in two 24-h food diaries on two separate days after the screening visit. As support, participants received a portion guide containing pictures of different portion sizes for all main food groups and an information leaflet with additional instructions. The portion guide and the food diaries were provided by the Swedish Food Agency (Livsmedelsverket, 2024). The 24-h recall and the two 24-h food diaries were used to evaluate if dietary fiber and protein intake were matching with the inclusion criteria. During the continuation of the study, three 24-h food diaries were collected each week on two weekdays and one weekend day of choice with at least one day in-between.

The GSRS was filled in by the participants on the evening before each visit to monitor gastrointestinal symptoms. The questionnaire includes 15 questions on symptoms during the past seven days based on a seven-point graded Likert-type scale (Kulich et al., 2008). Specifically, a score of 1 represents no discomfort at all, 2 minor discomfort, 3 mild discomfort, 4 moderate discomfort, 5 moderately severe discomfort, 6 severe discomfort, and 7 very severe discomfort. Additionally, participants were asked to document every bowel movement in a BSS diary regarding stool consistency and frequency. The BSS classifies stool consistency on a scale from 1 (hard stool) to 7 (liquid stool) as an estimation of gastrointestinal transit time (Lewis & Heaton, 1997). Furthermore, an IPAQ questionnaire was completed on the evening before each visit. The questionnaire covers different physical activity levels during the past seven days and was applied as a proxy of lifestyle maintenance (Craig et al., 2003).

### 2.7. Cell Culture

Human colonic epithelial cell caco-2 were cultured in Dulbecco’s Modified Eagle Medium (DMEM) containing penicillin and streptomycin with 10 % Fetal Bovine Serum (FBS) at 37 °C in a humidified atmosphere of 5 % CO_2_. Cells were passed to new culture plates by using trypsin/EDTA when they reached 80–90 % of confluence. Subsequently, cells were used to evaluate cytotoxicity of fecal water.

### 2.8. Resazurin Cell Viability and Lactate Dehydrogenase (LDH) Assays

Cells (1 × 10^4^ cells/well) were plated in a 96-well plate overnight, then 10 % (v/v) of the culture medium was replaced with filtered (0.22 µm) fecal water, which was resuspended in sterile phosphate buffered saline (PBS) at a 1:10 (w/v) ratio. A cell control had 10 % of DMEM medium replaced by sterile PBS, and a cell death control was treated with 0.02 % Triton X-100 (Sigma-Aldrich, Germany). After 24h of incubation, 20 µL of cell supernatant was transferred to measure extracellular LDH, while cells were added with 20 µL of resazurin dye solution (Sigma-Aldrich, Germany) and incubated for 3h.

For LDH determination, the commercially available LDH Activity Assay Kit was used (Sigma-Aldrich, Germany) according to the manufacturer’s instructions. Briefly, nicotinamide adenine dinucleotide hydrogen standards were used, and 50 µL of the master reaction mix was added to each well. The plate was mixed using a horizontal shaker protected from the light for 2 min and absorbance was measured at 450 nm in a microplate reader at 37 °C (Bio-Rad, USA), taking measurements every 5 min until the most active sample is greater than the highest standard. The final measurement was the one at which the most active sample was near to exceeding the highest standard, but which fell within the linear range of the standard curve. For the resazurin assay, fluorescence was monitored at a wavelength of 590 nm using an excitation wavelength of 560 nm (Bio-Rad, USA).

### 2.9. Power Calculation

The power calculation was based on the investigation of SCFA associated with protein fermentation in the gut. Prior research showed that dietary plant-based protein supplementation induced 8.42 % of increase in butyrate and a 7.54 % of increase in isovalerate (both expressed as relative proportions of total SCFA) over one week (Beaumont et al., 2017). The standard deviation of these differences was 7.54. For our study, we anticipated detecting at least 50 % of that effect size in a population with normal weight, using a Wilcox signed-rank test, 80 % power, and a 2-tailed significance level of 0.05. Our calculations indicated that a sample size of 32 subjects would detect a 0.53 standard deviation change in fecal SCFA.

### 2.10. Statistical Analyses

Week 1–3 was considered a training period for diary and questionnaire completion to ensure reliable data during week 4–8. Therefore, the first three weeks were not included in the final analysis. As collected questionnaire data included information of multiple days over a course of one week (24-h food diary), or whole weeks in general (GSRS, BSS and IPAQ) we refer to the timepoints as week rather than visit to highlight the different dimensions in the collected data.

In general, a repeated measures one-way analysis of variance (rmANOVA) for within group analyses and two-way rmANOVA for between group analyses was performed for normally distributed data after log-transformation. In case of missing values, a mixed-effects-analysis was used instead. Post-hoc analyses were conducted using the Dunnett’s and Šídák’s multiple comparisons test, respectively. In contrast, cell proliferation was analyzed using an ordinary one-way analysis of variance with Tukey’s multiple comparisons and differences in cytotoxicity were assessed with paired t-tests.

Non-normally distributed data was analyzed using the Friedman test for within group analyses, followed by the Dunn’s multiple comparison test. Between-group analyses were performed using the multiple Mann-Whitney test with Holm-Šídák multiple comparisons.

Preprocessing for gut microbiota composition analysis included agglomeration to species level, filtering Chordata sequences and transforming counts to relative counts per sample (per cent). Repeated measures correlation analyses were performed on log-transformed, scaled and sparsity-reduced data using Spearman’s rank correlation. The sparsity reduction removed operational taxonomic units with reads in the lower 30 % percentile and that appeared in less than two samples. Multiple comparisons were accounted for by using the Benjamini-Hochberg correction. Additionally, a random forest model (package randomForest; version 4.7-1.1) using package caret (version 6.0-94) for the optimization of the hyperparameter mtry was trained to classify before (visit 1–4) and under intervention samples (visit 5-8).

Statistical analyses were performed using GraphPad Prism Version 10.4.1 (GraphPad Software, USA) and R (version 4.3.2) using the package phyloseq (version 1.44.0). For all described analyses, results with a p-value of p<0.05 were considered statistically significant. Results are presented as mean ± SD for parametric data and median (IQR) for non-parametric data.

## 3. Results

A total of 311 people expressed interest in participating in the study, of whom 79 were screened, and 37 met eligible criteria. Following screening, six individuals were no longer interested in participating, and during the study, two participants withdrew due to personal reasons or a desire to discontinue the protein supplementation. Consequently, the study was completed with 29 participants (**Supplementary** Figure 1). Baseline characteristics of the study population are summarized in **Table 1**.

**Table 1.** Participant’s Baseline Characteristics at Visit 4.

|  |  | *Median (IQR)* | *n* |
| --- | --- | --- | --- |
| **Age (years)** |  |  |  |
|  | Total | 26 (22–33) | 29 |
|  | Female | 25 (21–29) | 20 |
|  | Male | 29 (24–37) | 9 |
| **Body Mass Index (kg/m<sup>2</sup>)** |  |  |  |
|  | Total | 23.00 (21.90–26.28) | 28 |
|  | Female | 22.50 (21.90–26.40) | 19 |
|  | Male | 24.30 (21.95–26.30) | 9 |

### 3.1. Protein Intake Compliance, Gastrointestinal Symptoms, Dietary and Lifestyle Habits

Neither energy nor dietary fiber intake changed significantly during the intervention. However, the intake of carbohydrates was significantly lower from week 7 onwards compared to week 4 (week 7: p=0.0002, week 8: p=0.0329), and fat intake was significantly reduced in week 8 (p=0.0013). As expected, protein intake increased significantly each week both when expressed as energy percentage (week 5: p=0.0004, week 6: p<0.0001, week 7: p<0.0001, week 8: p<0.0001) as well as per g/kg body mass/day (week 5: p=0.0026, week 6: p<0.0001, week 7: p<0.0001, week 8: p<0.0001). Protein intake measured as g/kg body mass/day, calculated from 24-hour urinary nitrogen also showed a significant increase from visit 7 onwards (visit 7: p=0.0056, visit 8: p=0.0114). Original data of all analyzed parameters are summarized in **Table 2**. As a proxy of compliance, 95.1 % of the empty protein supplements bags were returned by the participants.

**Table 2.** Dietary Intake According to 24-h Food Diaries and Protein Intake Calculated from Urea Nitrogen Obtained from 24-h Urine Samples.

|  | **Week 4** |  | **Week 5** |  | **Week 6** |  | **Week 7** |  | **Week 8** |  |
| --- | --- | --- | --- | --- | --- | --- | --- | --- | --- | --- |
|  | *Median (IQR)* | *n* | *Median (IQR)* | *n* | *Median (IQR)* | *n* | *Median (IQR)* | *n* | *Median (IQR)* | *n* |
| **Dietary Characteristics** |  |  |  |  |  |  |  |  |  |  |
| Energy (kcal/day) | 2104 (1735-2654) | 25 | 1974 (1757-2413) | 26 | 2160 (1945-2468) | 25 | 2017 (1933-2320) | 26 | 2195 (1941-2493) | 26 |
| Fiber (g/day) | 20.6 (16.5-25.1) | 25 | 22.5 (15.7-25.6) | 26 | 19.2 (16.4-22.4) | 25 | 19.0 (15.8-23.8) | 26 | 21.6 (17.2-24.0) | 26 |
| Carbohydrates (E %) | 45.4 (41.6-48.6) | 25 | 42.6 (38.5-48.5) | 26 | 41.5 (38.4-44.7) | 25 | 39.2 (37.5-42.2)*** | 26 | 40.1 (37.3-44.5)* | 26 |
| Fat (E %) | 38.5 (35.8-41.8) | 25 | 38.9 (35.5-42.4) | 26 | 37.1 (34.3-41.5) | 25 | 35.9 (31.0-39.7) | 26 | 34.3 (29.7-36.7)*** | 26 |
| Protein (E %) | 15.2 (13.6-17.8) | 25 | 17.3 (15.3-23.1)*** | 26 | 21.0 (18.0-24.2)**** | 25 | 23.2 (21.0-26.0)**** | 26 | 24.5 (22.4-27.0)**** | 26 |
| Protein (g/kg body mass/day) | 1.1 (0.9-1.2) | 25 | 1.3 (1.1-1.5)** | 26 | 1.6 (1.5-1.9)**** | 25 | 1.7 (1.6-1.9)**** | 26 | 2.0 (1.8-2.2)**** | 26 |
| **24-h Urea Nitrogen** |  |  |  |  |  |  |  |  |  |  |
|  | **Visit 4** |  | **Visit 5** |  | **Visit 6** |  | **Visit 7** |  | **Visit 8** |  |
| Protein (g/kg body mass/day) | 1.2 (0.8-1.6) | 16 | 1.6 (1.1-2.0) | 18 | 1.9 (1.3-2.2) | 13 | 1.6 (0.8-1.8)** | 12 | 2.0 (1.6-2.4)* | 17 |

Data is shown as median (IQR) and includes results from a mixed-effects analysis including Dunnett's multiple comparisons of each intervention visit/week vs. the baseline (visit/week 4); Statistical results are based on log-transformed data; \* p<0.05, \*\* p<0.01, \*\*\* p<0.001, \*\*\*\* p<0.0001; E %: energy percentage.

The median of the weekly reported BSS during the intervention period ranged from 3.8–4.3 which lies within the normal range of 3–5. The daily stool frequency during the intervention ranged from 0.9–1.1. There were no statistically significant differences in stool type and frequency during the intervention indicating no alterations due the increased protein intake (**Supplementary Table 2**). Since the study population is healthy, the average scores for the GSRS were low, as expected, with a median ranging from 1–2. Statistical analyses revealed no significant changes of the total GSRS score during the intervention (**Supplementary Table 2**). Physical activity levels measured in MET-min/week were statistically significant different (p=0.0031), however post-hoc multiple comparisons failed to detect significances. Physical activity, measured in physical activity levels (PAL) did not change significantly (**Supplementary Table 2**).

### 3.2. Surrogate Markers of Health

From visits 4–8, fasting blood, urine, and fecal samples were collected, and their correspondent biochemical measurements are summarized in **Table 3**. Plasma triglycerides were significantly decreased comparing visit 6 and 8 to visit 4 (p=0.0361), while plasma urea was significantly increased at visit 8 (p=0.0092). Plasma creatinine levels were significantly lower on all intervention visits (visit 5: p=0.0022, visit 6: p=0.0198, visit 7: p=0.0344, visit 8: p=0.0453). Despite limitations due to incomplete sample collection (53 % of completed samples), 24-h urine urea levels showed a significant increase by visit 8 (p=0.0494). In fecal samples, calprotectin levels increased significantly (visit 5: p=0.0194, visit 6: p=0.0.0236, visit 7: p=0.0.0236, visit 8: p=0.0021), whereas fecal ammonia levels decreased significantly (visit 5: p=0.0343, visit 7: p<0.0001, visit 8: p=0.0005). For SCFA, valerate significantly increased (visit 8: p=0.0315). Pearson correlation between ammonia in feces and urea in blood resulted in R^2^=0.8443, r=-0.9189 (p=0.0274).

**Table 3.**
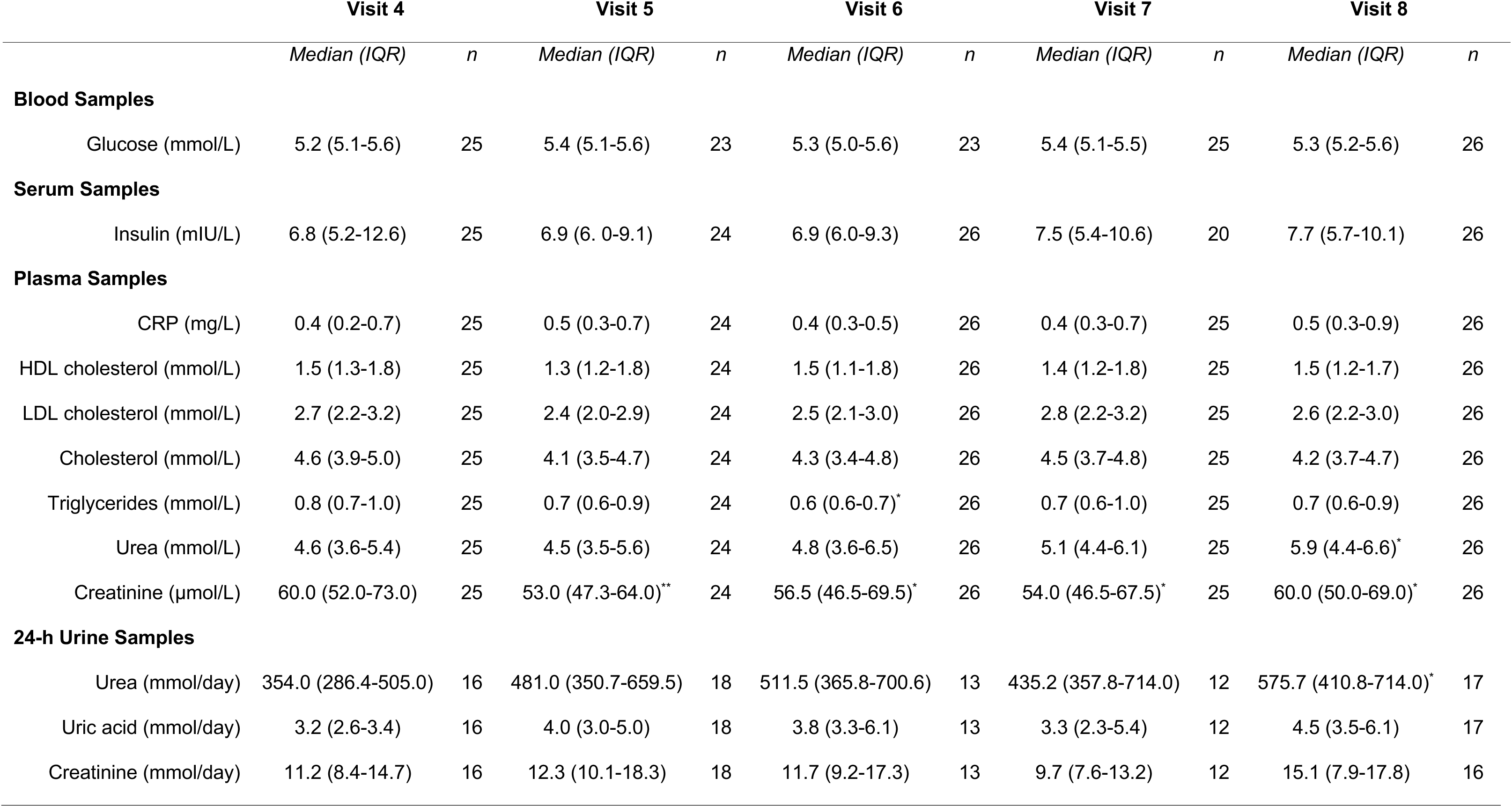
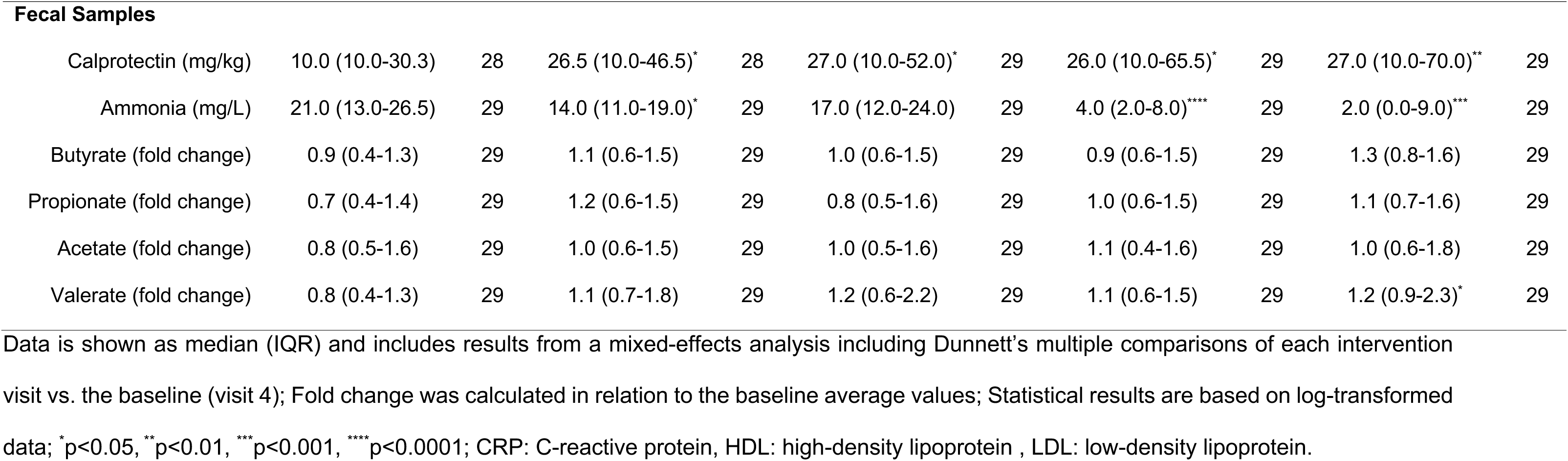
Biochemical Values Throughout the Intervention.

|  | Visit 4 |  | Visit 5 |  | Visit 6 |  | Visit 7 |  | Visit 8 |  |
| --- | --- | --- | --- | --- | --- | --- | --- | --- | --- | --- |
|  | *Median (IQR)* | *n* | *Median (IQR)* | *n* | *Median (IQR)* | *n* | *Median (IQR)* | *n* | *Median (IQR)* | *n* |
| **Blood Samples** |  |  |  |  |  |  |  |  |  |  |
| Glucose (mmol/L) | 5.2 (5.1-5.6) | 25 | 5.4 (5.1-5.6) | 23 | 5.3 (5.0-5.6) | 23 | 5.4 (5.1-5.5) | 25 | 5.3 (5.2-5.6) | 26 |
| **Serum Samples** |  |  |  |  |  |  |  |  |  |  |
| Insulin (mIU/L) | 6.8 (5.2-12.6) | 25 | 6.9 (6. 0-9.1) | 24 | 6.9 (6.0-9.3) | 26 | 7.5 (5.4-10.6) | 20 | 7.7 (5.7-10.1) | 26 |
| **Plasma Samples** |  |  |  |  |  |  |  |  |  |  |
| CRP (mg/L) | 0.4 (0.2-0.7) | 25 | 0.5 (0.3-0.7) | 24 | 0.4 (0.3-0.5) | 26 | 0.4 (0.3-0.7) | 25 | 0.5 (0.3-0.9) | 26 |
| HDL cholesterol (mmol/L) | 1.5 (1.3-1.8) | 25 | 1.3 (1.2-1.8) | 24 | 1.5 (1.1-1.8) | 26 | 1.4 (1.2-1.8) | 25 | 1.5 (1.2-1.7) | 26 |
| LDL cholesterol (mmol/L) | 2.7 (2.2-3.2) | 25 | 2.4 (2.0-2.9) | 24 | 2.5 (2.1-3.0) | 26 | 2.8 (2.2-3.2) | 25 | 2.6 (2.2-3.0) | 26 |
| Cholesterol (mmol/L) | 4.6 (3.9-5.0) | 25 | 4.1 (3.5-4.7) | 24 | 4.3 (3.4-4.8) | 26 | 4.5 (3.7-4.8) | 25 | 4.2 (3.7-4.7) | 26 |
| Triglycerides (mmol/L) | 0.8 (0.7-1.0) | 25 | 0.7 (0.6-0.9) | 24 | 0.6 (0.6-0.7)* | 26 | 0.7 (0.6-1.0) | 25 | 0.7 (0.6-0.9) | 26 |
| Urea (mmol/L) | 4.6 (3.6-5.4) | 25 | 4.5 (3.5-5.6) | 24 | 4.8 (3.6-6.5) | 26 | 5.1 (4.4-6.1) | 25 | 5.9 (4.4-6.6)* | 26 |
| Creatinine (μmol/L) | 60.0 (52.0-73.0) | 25 | 53.0 (47.3-64.0)** | 24 | 56.5 (46.5-69.5)* | 26 | 54.0 (46.5-67.5)* | 25 | 60.0 (50.0-69.0)* | 26 |
| **24-h Urine Samples** |  |  |  |  |  |  |  |  |  |  |
| Urea (mmol/day) | 354.0 (286.4-505.0) | 16 | 481.0 (350.7-659.5) | 18 | 511.5 (365.8-700.6) | 13 | 435.2 (357.8-714.0) | 12 | 575.7 (410.8-714.0)* | 17 |
| Uric acid (mmol/day) | 3.2 (2.6-3.4) | 16 | 4.0 (3.0-5.0) | 18 | 3.8 (3.3-6.1) | 13 | 3.3 (2.3-5.4) | 12 | 4.5 (3.5-6.1) | 17 |
| Creatinine (mmol/day) | 11.2 (8.4-14.7) | 16 | 12.3 (10.1-18.3) | 18 | 11.7 (9.2-17.3) | 13 | 9.7 (7.6-13.2) | 12 | 15.1 (7.9-17.8) | 16 |

| Fecal Samples |  |  |  |  |  |  |  |  |  |  |
| --- | --- | --- | --- | --- | --- | --- | --- | --- | --- | --- |
| Calprotectin (mg/kg) | 10.0 (10.0-30.3) | 28 | 26.5 (10.0-46.5)* | 28 | 27.0 (10.0-52.0)* | 29 | 26.0 (10.0-65.5)* | 29 | 27.0 (10.0-70.0)** | 29 |
| Ammonia (mg/L) | 21.0 (13.0-26.5) | 29 | 14.0 (11.0-19.0)* | 29 | 17.0 (12.0-24.0) | 29 | 4.0 (2.0-8.0)**** | 29 | 2.0 (0.0-9.0)*** | 29 |
| Butyrate (fold change) | 0.9 (0.4-1.3) | 29 | 1.1 (0.6-1.5) | 29 | 1.0 (0.6-1.5) | 29 | 0.9 (0.6-1.5) | 29 | 1.3 (0.8-1.6) | 29 |
| Propionate (fold change) | 0.7 (0.4-1.4) | 29 | 1.2 (0.6-1.5) | 29 | 0.8 (0.5-1.6) | 29 | 1.0 (0.6-1.5) | 29 | 1.1 (0.7-1.6) | 29 |
| Acetate (fold change) | 0.8 (0.5-1.6) | 29 | 1.0 (0.6-1.5) | 29 | 1.0 (0.5-1.6) | 29 | 1.1 (0.4-1.6) | 29 | 1.0 (0.6-1.8) | 29 |
| Valerate (fold change) | 0.8 (0.4-1.3) | 29 | 1.1 (0.7-1.8) | 29 | 1.2 (0.6-2.2) | 29 | 1.1 (0.6-1.5) | 29 | 1.2 (0.9-2.3)* | 29 |

Data is shown as median (IQR) and includes results from a mixed-effects analysis including Dunnett's multiple comparisons of each intervention visit vs. the baseline (visit 4); Fold change was calculated in relation to the baseline average values; Statistical results are based on log-transformed data; \*p<0.05, \*\*p<0.01, \*\*\*p<0.001, \*\*\*\*p<0.0001; CRP: C-reactive protein, HDL: high-density lipoprotein , LDL: low-density lipoprotein.

### 3.3. Subgrouping by Fecal Calprotectin

Interestingly, together with a significant increase in fecal calprotectin over time during the intervention, the variability in calprotectin levels among participants also increased. To investigate whether this variability was driven by a subset of participants, we divided the group based on fecal calprotectin levels at visit 8, splitting participants at the median into a high-calprotectin (HC) and a low-calprotectin (LC) subgroup for further analysis. These groups were not significantly different regarding age, sex, BMI and their dietary fiber intake (results not shown). **Figure 2A** illustrates that calprotectin increased only in the HC subgroup significantly compared with visit 4 (visit 6: p=0.0367, visit 7: p=0.0320, visit 8: p<0.0001). No significant changes were observed in the LC subgroup. The subgroups differed significantly starting from visit 6 onwards (visit 6: p=0.0075, visit 7: p=0.0051, visit 8: p<0.0001). The baseline comparison at visit 4 showed a trend for significance (p=0.0706). Further analysis revealed a general trend of increasing SCFA levels over time for LC subgroup (**Figure 2B**), and levels of valerate and butyrate were significantly higher at visit 8 (p=0.0365 and p=0.0427, respectively). No significant changes were observed in the HC subgroup nor between LC and HC subgroups.

**Figure 2.**
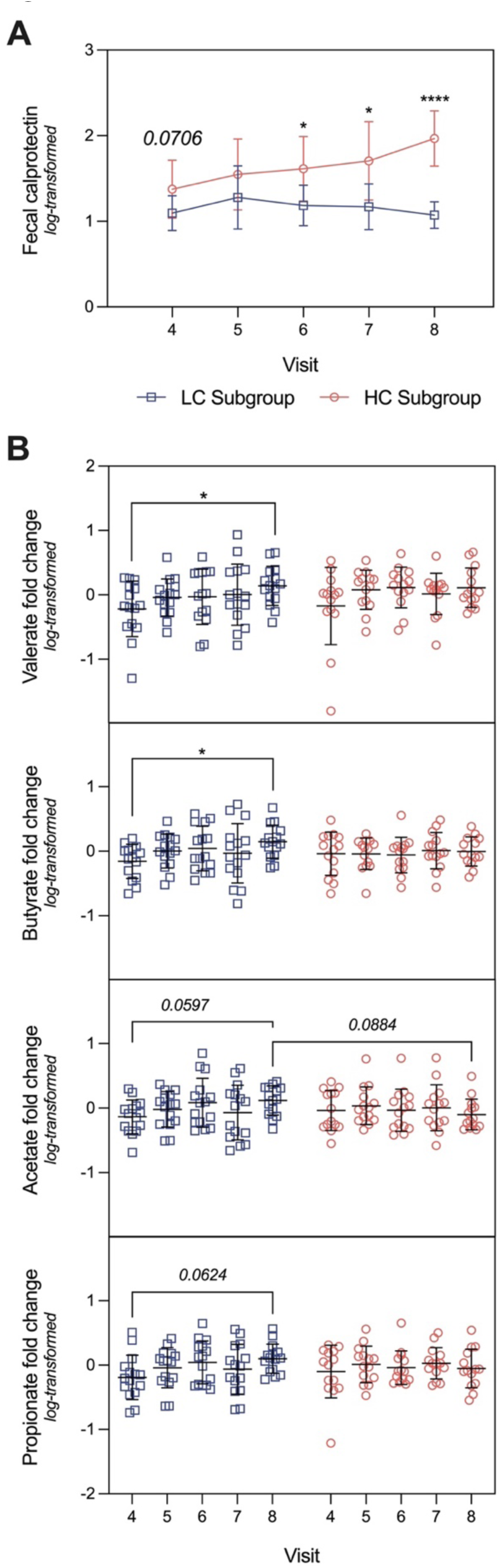
Participants Divided by Fecal Protein Response at Visit 8. A) Differences in fecal calprotectin levels between participant subgroups split by upper and lower 50 % percentile based on visit 8, including results from a mixed-effects analysis with Šídák’s multiple comparisons test. B) Fecal butyrate, propionate, acetate and valerate fold change over the intervention in HC and LC subgroups, including results from a two-way ANOVA with Šídák’s multiple comparisons test. Statistical results are based on log-transformed data; *p<0.05, ****p<0.0001. All results with p<0.1 are included in the figure. LC: low-calprotectin (n=15), HC: high-calprotectin (n=14), ANOVA: analysis of variance.

### 3.4. Colonic Cell Viability and Cytotoxicity

Cell proliferation was not differently affected by the fecal water from baseline (visit 4) and from the intervention’s end (visit 8). Additionally, no differences were observed comparing fecal water samples to the control cells only treated with cell culture medium (**Figure 3A**).

**Figure 3.**
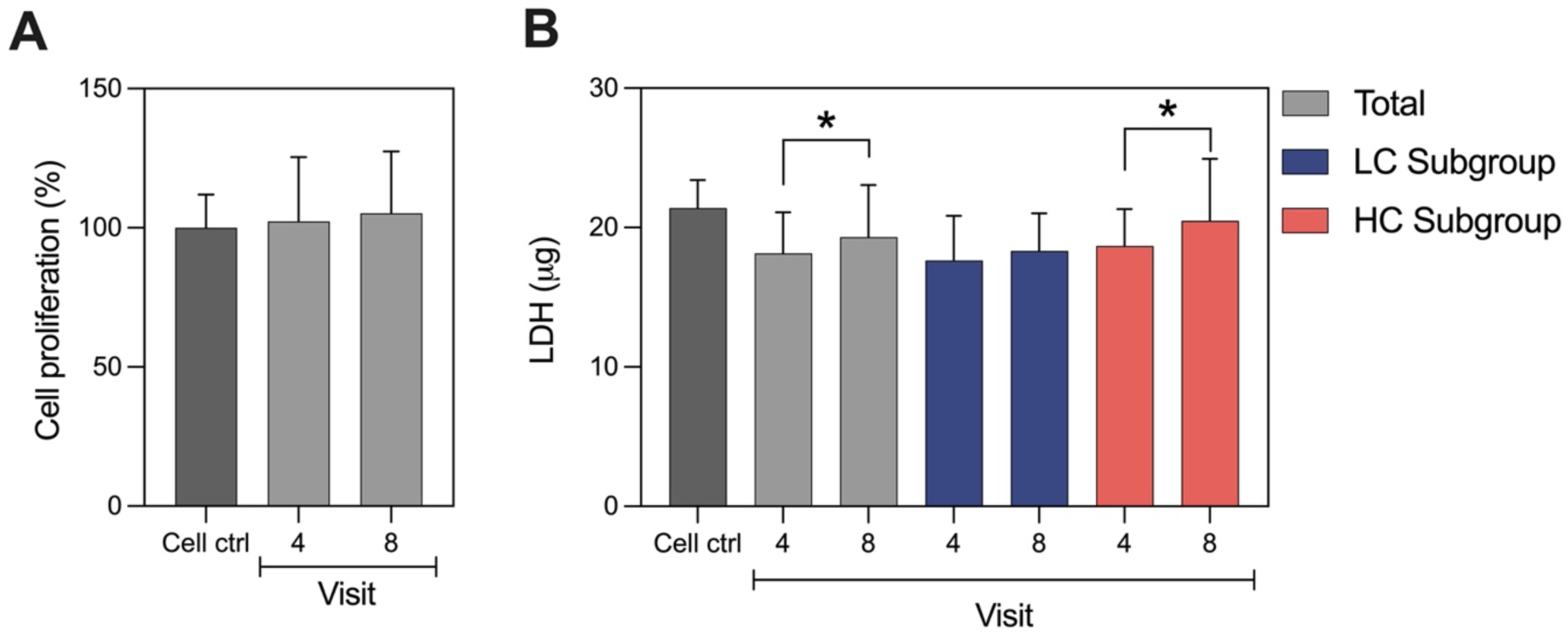
Cell Proliferation and Cytotoxicity when Treated with Fecal Water. A) Cell proliferation assessed using the resazurin method. B) LDH release in the total population and in each subgroup (HC and LC) as a measure of cytotoxicity including results from paired t-tests. *p<0.05. LC: low-calprotectin (n=15), HC: high-calprotectin (n=14), LDH: lactate dehydrogenase.

To further assess cytotoxicity, we measured LDH, a cytosolic enzyme released from damaged cells. Extracellular LDH levels increased significantly at visit 8 compared to visit 4 (p=0.0100). However, no differences were observed comparing fecal water to the control cells. Since calprotectin can promote cytotoxicity in epithelial cells (Shabani et al., 2020), we investigated whether fecal water would exert distinct cytotoxic effects in LC and HC subgroups. The HC subgroup exhibited higher LDH release after the intervention (p=0.0077), while no change was observed for the LC subgroup between visits 4 and 8 (**Figure 3B**).

### 3.5. Microbiota Composition

The average read length and depth per sample was 144 bp and 13.9 Mio, respectively. There was no significant difference in α and β diversity as measured by the Shannon-Diversity Index and the Bray-Curtis Index during the intervention and within LC and HC subgroups (**Figure 4A**). The most abundant phylum among the study population was Firmicutes, followed by Bacteroidetes, Actinobacteria and Verrucomicrobiota (**Figure 4B**).

**Figure 4.**
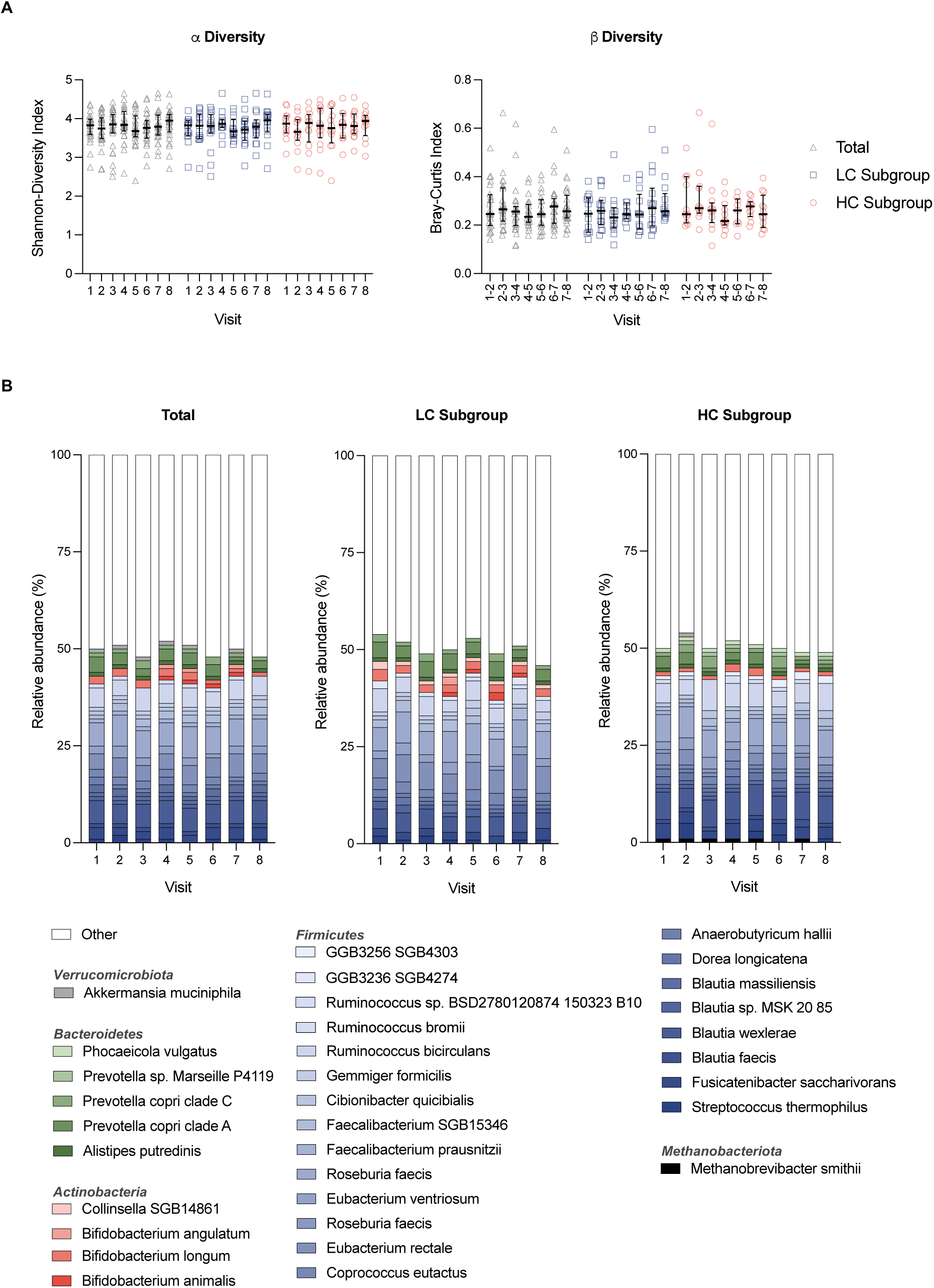
Changes in α and β Diversity and Microbiota Composition. A) Individual values for Alpha-diversity, Shannon-Diversity Index, and Beta-diversity, Bray-Curtis Index, for the total population as well as both subgroups, including median (IQR). B) Top 15 identified species in all samples belonging to each respective week. LC: low-calprotectin (n=15), HC: high-calprotectin (n=14).

To investigate if the intervention has led to changes in species abundances, we analyzed if relative abundances correlate with visits 4–8. We found a significant and positive correlation for the species *L. frumenti* (correlation coefficient: 0.55, p<0.0001), *O. splanchnicus* (correlation coefficient: 0.53, p<0.001), and *L. crispatus* (correlation coefficient: 0.43, p=0.002). In contrast, relative abundances of *B. longum* and *B. catenulatum* were significantly negatively correlated with the intervention visits (correlation coefficient: -0.39, p=0.010 and correlation coefficient: -0.37, p=0.019, respectively). If participants were grouped into LC and HC, a significant positive correlation remained for *L. frumenti* (correlation coefficient: 0.59, p=0.003), *O. splanchnicus* (correlation coefficient: 0.59, p=0.003) and *L. crispatus* (correlation coefficient: 0.57, p=0.004) in the HC subgroup. Subsequently, we analyzed relative abundances of the correlating species to further explore potential differences (**Figure 5**). In the total population, significantly higher abundances of *L. frumenti* were present at visit 5–8 compared to visit 4 (visit 5: p<0.0001, visit 6: p<0.0001, visit 7: p<0.0001, visit 8: p<0.0001). This was also observed in the LC subgroup (visit 5: p=0.0064, visit 6: p=0.0014, visit 7: p=0.0012, visit 8: p=0.0095) and in the HC subgroup (visit 5: p=0.0208, visit 6: p=0.0020, visit 7: p=0.0042, visit 8: p=0.0001). Relative abundances of *O. splanchnicus* increased significantly at visit 7 and 8 (visit 7: p=0.0144, visit 8: p=0.0002) in the total population. Whereas the observed differences at visit 7 and 8 were maintained for HC (visit 7: p=0.0383, visit 8: p=0.0141), only differences at visit 8 remained significant within the LC subgroup (p=0.0288). For *L. crispatus,* results showed a significantly increased abundance at visits 6-8 (visit 6: p=0.0056, visit 7: p=0.0038, visit 8: p=0.0012) in the total population. Within the subgroups we observed significantly higher abundances at visit 6 for LC (p=0.0363) and visit 8 for HC (p= 0.0162). In contrast to the previously mentioned species, relative abundances of *B. longum* were significantly lower at visit 6 and visit 8 in the total population (visit 6: p=0.0131, visit 8: p=0.0006). Subgroup analyses revealed lower abundances at visit 8 in the LC subgroup (p=0.0157) and lower abundances at visit 6 in the HC subgroup (p=0.0027). For *B. catenulatum*, the Friedman tests was significant for the total population (p=0.0004) and the LC subgroup (p=0.0063), however, multiple comparisons failed to detect significances. Within the HC subgroup, none of the conducted tests were significant.

**Figure 5.**
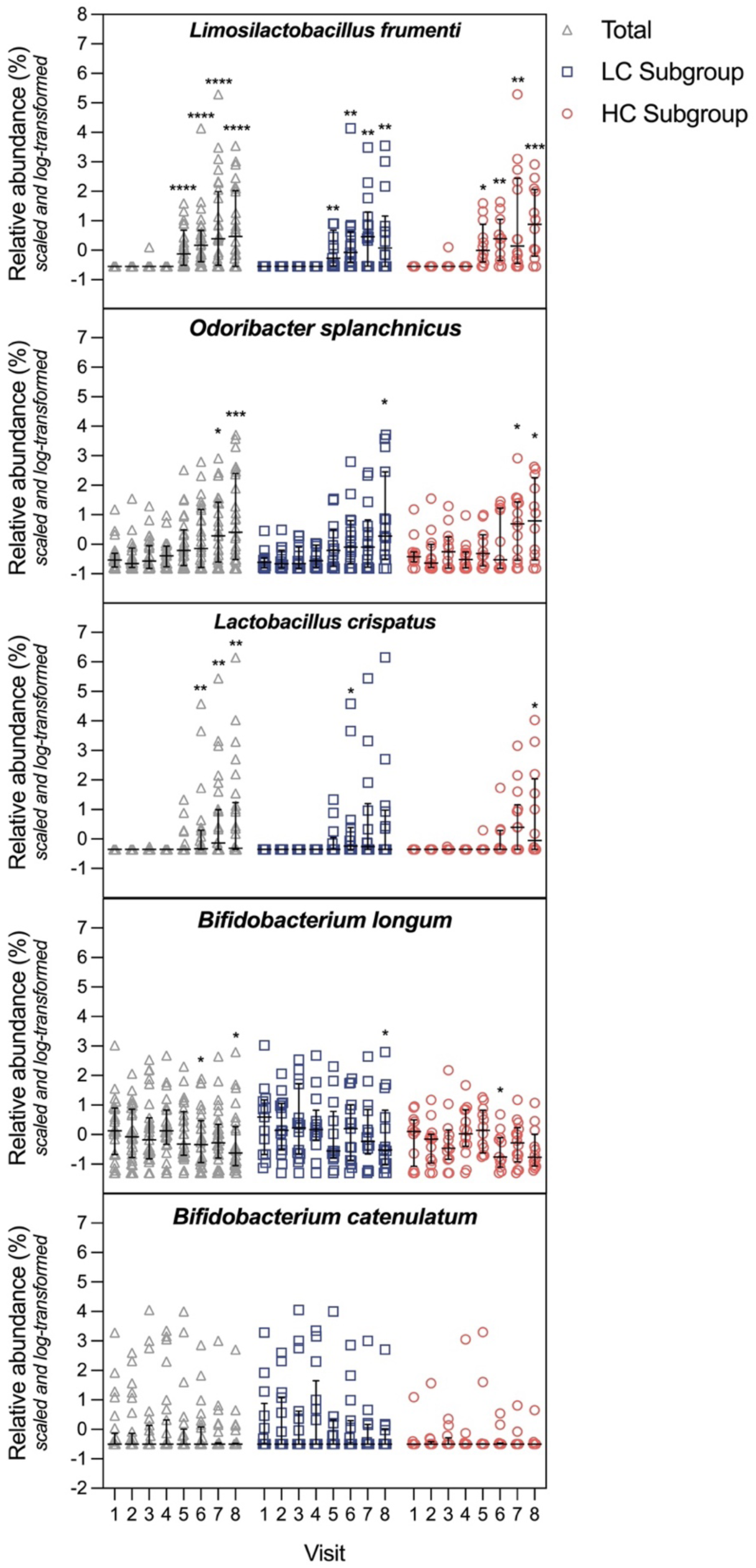
Relative abundances of species correlating with the intervention visits. Individual relative abundance values for *L. frumenti*, *O. splanchnicus*, *L. crispatus*, *B. longum* as well as *B. catenulatum* for the total population as well as both subgroups (HC and LC). The figures include median (IQR) and results from the Friedman test with Dunn’s post-hoc test comparing visits 5–8 to visit 4; *p<0.05, **p<0.01, ***p<0.001, ****p<0.0001; LC: low-calprotectin (n=15), HC: high-calprotectin (n=14).

There were no significant differences in relative abundances of the correlating species between the subgroups.

The random forest model used to identify important variables confirmed results from correlation analysis, with *L. frumenti* as the by far most important variable (permutation p-value of 0 in 200 repetitions). Subsequently, we investigated if the six relevant species also correlate with any other biological markers in the intervention study. In the total population, relative abundances of *L. frumenti* significantly correlated with fecal ammonia (correlation coefficient: -0.41, p=0.001), plasma urea (correlation coefficient: 0.37, p=0.009) and fecal calprotectin (correlation coefficient: 0.33, p=0.009). *O. splanchnicus* significantly correlated with fecal ammonia (correlation coefficient: -0.34, p=0.009) and *L. crispatus* significantly correlated with fecal ammonia (correlation coefficient: -0.33, p=0.009) as well as plasma urea (correlation coefficient: 0.31, p=0.025). Furthermore, we found a significant correlation between relative abundances of *B. longum* and plasma cholesterol (correlation coefficient: -0.32, p=0.025). In the HC subgroup, *L. frumenti* and *L. crispatus* significantly correlated with fecal calprotectin (correlation coefficient: 0.57, p=0.0007 and correlation coefficient: 0.45, p=0.013, respectively) as well as fecal ammonia (correlation coefficient: -0.45, p=0.013 and correlation coefficient: - 0.46, p=0.013, respectively).

## 4. Discussion

This is the first dietary intervention to demonstrate a significant increase in fecal calprotectin levels in a subset of healthy participants in response to increasing amounts of isolated pea protein supplementation. Additionally, fecal butyrate and valerate levels were significantly elevated in participants with stable fecal calprotectin. These findings highlight the importance of assessing plant-based protein intake in healthy individuals to establish potential links with disease risk in the future. Furthermore, this study extends the existing literature on generalized dietary effects by providing insights into which biomarkers exhibit interindividual variability under comparable dietary conditions.

### 4.1. Effects of Protein Supplementation on Biomarkers of Gut Health and Microbiota Composition

Fecal calprotectin is a non-specific but sensitive marker of intestinal inflammation, released by neutrophils in the intestinal mucosa during inflammatory processes and commonly used to monitor inflammatory bowel disease (Bressler et al., 2015; Kapel et al., 2023). The observed increase in fecal calprotectin levels in certain individuals points towards the presence of responders and non-responders. The differences between subgroup baseline values of calprotectin showed a trend towards significance, suggesting that an increased intake of plant-based proteins may promote inflammation in predisposed individuals (Mendall et al., 2016). In line with previous research, higher LDH cytotoxicity of fecal water was only observed in the

HC subgroup (Shabani et al., 2020), highlighting that LDH cytotoxicity may be linked to increased inflammation (Kämpfer et al., 2017). Despite significant differences between HC and LC, most of the observed fecal calprotectin values were within the physiological range and below the 80 mg/kg cut-off applied in the used assay.

Interestingly, the rather detrimental metabolite ammonia, resulting from proteolytic fermentation, decreased in feces with an increase in protein intake. This is aligned with another study showing that labeled nitrogen excretion in feces tended to be lower in a high-protein diet compared with low-protein diet (Windey et al., 2012b). Ammonia can be used by the gut bacteria or be absorbed by the colonic mucosa and converted in the liver to urea, being excreted in the urine later (Wrong & Vince, 1984). Such metabolism goes in hand with the negative correlation between fecal ammonia and plasma urea, indicating an increased availability of ammonia in the gut to being taken up.

Another explanation for the decrease of fecal ammonia is its utilization by certain microbial taxa in the gut as a nitrogen source for generating microbial biomass (Bergen & Wu, 2009). The observed increase of relative abundance of *L. frumenti*, *L. crispatus, and O. splanchnicus* and their negative correlations with fecal ammonia may strengthen this hypothesis. While literature on underlying metabolic pathways of the specific strains is scarce, research on lactic acid bacteria has described their proteolytic properties and their growth dependence on nitrogen, even though the extent of utilization can differ between strains and the nitrogen source (Hu et al., 2024; M. Liu et al., 2010). Species within the Genus *Lactobacillus* have been linked to beneficial effects in the past, especially for their probiotic characteristics (Dempsey & Corr, 2022; Shah et al., 2024). Probiotics are “live microorganisms which when administered in adequate amounts, confer a health benefit on the host” (Hill et al., 2014), for instance by promoting intestinal barrier integrity and function, improving insulin sensitivity and exerting immunomodulatory effects (Gul & Durante-Mangoni, 2024). However, research on the role specific species mentioned above in human health is still sparse.

Valerate is strongly linked to proteolytic fermentation (Neis et al., 2015) and its increase was observed in overweight adults receiving soy protein supplementation compared to control or casein groups (Beaumont et al., 2017), aligning with our findings. In our study, only the LC subgroup showed increased butyrate, with a tendency for higher propionate and acetate. Although bacteria can produce butyrate from protein, in human studies only propionate, and not butyrate, has been associated with higher protein intake (Mak et al., 2025). In *in vitro* fermentations, the protein-to-fiber ratio appears to be a key factor (Jackson et al., 2024). Notably, despite similar protein-to-fiber ratios, the LC and HC subgroups exhibited different butyrate trends, suggesting individual variability in response.

Although there is limited evidence in the literature on the association between SCFA and fecal calprotectin, fecal calprotectin levels are strongly correlated with intestinal mucosal inflammation where levels of SCFA levels play a key role in immune regulation (Jukic et al., 2021; X. Liu et al., 2023; Parada Venegas et al., 2019). Taken together, these findings point towards a potential link between SCFA-mediated modulation of gut inflammation and lower calprotectin levels.

While markers typically associated with gut health - such as higher SCFA levels and lower calprotectin - were observed only in a subset of participants, all individuals exhibited an increase in some beneficial bacteria. Such pattern was seen following the increased consumption of isolated pea protein despite unchanged dietary fiber intake during the intervention and between LC and HC subgroups. These results indicate that observed changes in relative abundances and SCFA levels may not result from altered fermentation of indigestible carbohydrates. However, the presence of other phytochemicals in the protein isolate such as flavonoids (Cosson et al., 2022), may have impacted the observed changes in relative abundance, as they have been shown to affect gut microbiota composition (Mithul Aravind et al., 2021; Zhou et al., 2024).

This exploratory study suggests the presence of both species-dependent and species-independent microbiota-driven mechanisms triggered by dietary components, highlighting the need for further investigation into microbiota functionality. The observed changes in species abundance, SCFA, and calprotectin indicate that unabsorbed protein isolate reached the colon.

### 4.2. Strengths & Limitations

The combination of self-reported and several objective measurements of protein intake compliance (24-h food diaries, returned protein bags, as well as blood and 24-h urine urea levels) suggests high adherence to the intervention. Additionally, participants maintained an isocaloric diet by slightly decreasing their carbohydrate and fat intake, which remained close to or within the Nordic Nutrition Recommendations for a normal diet (45–60 E % and 25–40 E%, respectively) (Blomhoff et al., 2023). Still, this could limit the ability to attribute all observed effects solely to pea protein isolate. To ensure a homogeneous study population, we included only participants within predefined dietary fiber and protein intake ranges, further strengthening the observed results.

We concluded the study with 29 participants, three less than anticipated. However, we decided not to proceed with further recruitment, a decision influenced by the ongoing COVID-19 pandemic and logistical constraints. Additionally, our subgroup analyses based on fecal calprotectin were of exploratory nature, wherefore they may lack sufficient sample size. In addition, the predominance of young female participants limits the generalizability of our findings to other populations and the assessment of sex-related differences.

The weekly increment in protein intake was designed to assess the feasibility of protein supplementation and to determine the threshold for metabolic changes. However, this approach limits our ability to distinguish whether the observed effects were driven by intervention duration or by the increasing protein dose. Nevertheless, the gradual increase facilitated compliance and adaptation while allowing participants to reach a high-protein intake without drastic dietary shifts. The absence of a control group also makes it challenging to rule out potential reactivity effects due to sample and questionnaire collection. However, the inclusion of a 4-week baseline period before the intervention likely mitigated such biases, as participants had time to adapt to study procedures before the intervention began.

## 5. Conclusion

As societal shifts in protein intake influence both its quantity and source, it is essential to investigate the resulting impact on health biomarkers and gut microbiota. Our findings suggest that increasing doses of isolated pea protein can modulate the growth of beneficial bacteria and affect fecal calprotectin and SCFA levels in a subset of healthy participants. Lower calprotectin levels were associated with a tendency towards increased fecal SCFA, while participants with higher calprotectin showed only minimal SCFA changes and increased fecal water cytotoxicity *in vitro*. These results highlight the importance of assessing health markers related to plant-based protein intake in healthy populations, as well as the search for biological differential responses. Future research should include larger cohorts and control groups to further explore subgroup differences in gut inflammation.

## Author Contribution

S.B.R.P.: Conceptualization; Data curation; Formal analysis; Funding acquisition; Investigation; Methodology; Project administration; Resources; Supervision; Validation; Visualization; Writing - original draft; Writing - review & editing. A.K.: Data curation; Formal analysis; Investigation; Methodology; Supervision; Validation; Visualization; Writing - original draft; Writing - review & editing. K.D.: Formal analysis; Methodology; Visualization; Writing - review & editing. I.K. Data curation; Methodology; Writing - review & editing. V.C.C.A.: Formal analysis; Methodology; Visualization; Writing - review & editing. T.H.: Methodology; Writing - review & editing. M.L.: Conceptualization; Methodology; Writing - review & editing. D.R.: Formal analysis; Methodology; Visualization; Writing - review & editing. T.M.M.: Conceptualization; Project administration; Supervision; Validation. R.J.B.: Conceptualization; Funding acquisition; Investigation; Project administration; Resources; Supervision; Validation; Writing - review & editing. All authors had final approval of the submitted manuscript and published versions.

## Acknowledgments

We thank all participants, the interns Sofie Raemisch and Giulia Bevilacqua, the study nurses who assisted in this dietary intervention and Lantmännen for providing the isolated pea protein used in the study. Furthermore, we would also like to acknowledge Clinical Genomics Örebro, Science for Life Laboratory, for providing expertise and service with sequencing. This work was supported by Örebro University Food and Health (ORU 1.4.1-00125/2021) and FORMAS (2020-02843 and 2021-02037). The DAAD and ERASMUS programs supported interns.

## References

Abe-Inge, V., Aidoo, R., Moncada de la Fuente, M., & Kwofie, E. M. (2024). Plant-based dietary shift: Current trends, barriers, and carriers. Trends in Food Science & Technology, 143, 104292. 10.1016/j.tifs.2023.104292

Alvarenga, L., Kemp, J. A., Baptista, B. G., Ribeiro, M., Lima, L. S., & Mafra, D. (2024). Production of Toxins by the Gut Microbiota: The Role of Dietary Protein. Current Nutrition Reports, 13(2), 340–350. 10.1007/s13668-024-00535-x

Ashkar, F., & Wu, J. (2023). Effects of Food Factors and Processing on Protein Digestibility and Gut Microbiota. Journal of Agricultural and Food Chemistry, 71(23), 8685–8698. 10.1021/acs.jafc.3c00442

Auer, J., Alminger, M., Marinea, M., Johansson, M., Zamaratskaia, G., Högberg, A., & Langton, M. (2024). Assessing the digestibility and estimated bioavailability/ bioaccessibility of plant-based proteins and minerals from soy, pea, and faba bean ingredients. LWT, 197, 115893. 10.1016/j.lwt.2024.115893

Beaumont, M., Portune, K. J., Steuer, N., Lan, A., Cerrudo, V., Audebert, M., Dumont, F., Mancano, G., Khodorova, N., Andriamihaja, M., Airinei, G., Tomé, D., Benamouzig, R., Davila, A.-M., Claus, S. P., Sanz, Y., & Blachier, F. (2017). Quantity and source of dietary protein influence metabolite production by gut microbiota and rectal mucosa gene expression: a randomized, parallel, double-blind trial in overweight humans. The American Journal of Clinical Nutrition, 106(4), 1005–1019. 10.3945/ajcn.117.158816

Benković, M., Jurinjak Tušek, A., Sokač Cvetnić, T., Jurina, T., Valinger, D., & Gajdoš Kljusurić, J. (2023). An Overview of Ingredients Used for Plant-Based Meat Analogue Production and Their Influence on Structural and Textural Properties of the Final Product. Gels, 9(12), 921. 10.3390/gels9120921

Bergen, W. G., & Wu, G. (2009). Intestinal Nitrogen Recycling and Utilization in Health and Disease. The Journal of Nutrition, 139(5), 821–825. 10.3945/jn.109.104497

Blachier, F., Mariotti, F., Huneau, J. F., & Tomé, D. (2007). Effects of amino acid-derived luminal metabolites on the colonic epithelium and physiopathological consequences. Amino Acids, 33(4), 547–562. 10.1007/s00726-006-0477-9

Blomhoff, R., Andersen, R., Arnesen, E. K., Christensen, J. J., Eneroth, H., Erkkola, M., Gudanaviciene, I., Halldórsson, Þ. I., Høyer-Lund, A., Lemming, E. W., Meltzer, H. M., Pitsi, T., Siksna, I., Þórsdóttir, I., & Trolle, E. (2023). Nordic Nutrition Recommendations 2023. 10.6027/nord2023-003

Bolte, L. A., Vich Vila, A., Imhann, F., Collij, V., Gacesa, R., Peters, V., Wijmenga, C., Kurilshikov, A., Campmans-Kuijpers, M. J. E., Fu, J., Dijkstra, G., Zhernakova, A., & Weersma, R. K. (2021). Long-term dietary patterns are associated with pro-inflammatory and anti-inflammatory features of the gut microbiome. Gut, 70(7), 1287–1298. 10.1136/gutjnl-2020-322670

Boven, L., Akkerman, R., & de Vos, P. (2024). Sustainable diets with plant-based proteins require considerations for prevention of proteolytic fermentation. Critical Reviews in Food Science and Nutrition, 1–11. 10.1080/10408398.2024.2352523

Bressler, B., Panaccione, R., Fedorak, R. N., & Seidman, E. G. (2015). Clinicians’ guide to the use of fecal calprotectin to identify and monitor disease activity in inflammatory bowel disease. Canadian Journal of Gastroenterology & Hepatology, 29(7), 369–372. 10.1155/2015/852723

Broucke, K., Duquenne, B., De Witte, B., De Paepe, E., De Smet, S., & Van Royen, G. (2025). Effect of salt, phosphate and protein content on the techno-functionality of plant-based proteins for hybrid meat product formulation. European Food Research and Technology. 10.1007/s00217-025-04692-3

Campos-Perez, W., & Martinez-Lopez, E. (2021). Effects of short chain fatty acids on metabolic and inflammatory processes in human health. Biochimica et Biophysica Acta (BBA) - Molecular and Cell Biology of Lipids, 1866(5), 158900. 10.1016/J.BBALIP.2021.158900

Cosson, A., Meudec, E., Ginies, C., Danel, A., Lieben, P., Descamps, N., Cheynier, V., Saint-Eve, A., & Souchon, I. (2022). Identification and quantification of key phytochemicals in peas – Linking compounds with sensory attributes. Food Chemistry, 385, 132615. 10.1016/j.foodchem.2022.132615

Craig, C. L., Marshall, A. L., Sjöström, M., Bauman, A. E., Booth, M. L., Ainsworth, B. E., Pratt, M., Ekelund, U., Yngve, A., Sallis, J. F., & Oja, P. (2003). International Physical Activity Questionnaire: 12-Country Reliability and Validity. Medicine & Science in Sports & Exercise, 35(8), 1381–1395. 10.1249/01.MSS.0000078924.61453.FB

Czaja, T. P., Beldring, S. N., Renaud, C., & Engelsen, S. B. (2025). Mimicking the properties of commercial chocolate mousses using plant proteins as foaming stabilisers. Texture, rheology, color and proton mobility. Food Research International, 212, 116450. 10.1016/j.foodres.2025.116450

David, L. A., Maurice, C. F., Carmody, R. N., Gootenberg, D. B., Button, J. E., Wolfe, B. E., Ling, A. V., Devlin, A. S., Varma, Y., Fischbach, M. A., Biddinger, S. B., Dutton, R. J., & Turnbaugh, P. J. (2014). Diet rapidly and reproducibly alters the human gut microbiome. Nature, 505(7484), 559–563. 10.1038/nature12820

Dempsey, E., & Corr, S. C. (2022). Lactobacillus spp. for Gastrointestinal Health: Current and Future Perspectives. Frontiers in Immunology, 13. 10.3389/fimmu.2022.840245

Ebert, S., Baune, M.-C., Broucke, K., Royen, G. Van, Terjung, N., Gibis, M., & Weiss, J. (2021). Buffering capacity of wet texturized plant proteins in comparison to pork meat. Food Research International, 150, 110803. 10.1016/j.foodres.2021.110803

Fung, K., Ooi, C., Zucker, M., Lockett, T., Williams, D., Cosgrove, L., & Topping, D. (2013). Colorectal Carcinogenesis: A Cellular Response to Sustained Risk Environment. International Journal of Molecular Sciences, 14(7), 13525–13541. 10.3390/ijms140713525

Gul, S., & Durante-Mangoni, E. (2024). Unraveling the Puzzle: Health Benefits of Probiotics—A Comprehensive Review. Journal of Clinical Medicine, 13(5), 1436. 10.3390/jcm13051436

Hill, C., Guarner, F., Reid, G., Gibson, G. R., Merenstein, D. J., Pot, B., Morelli, L., Canani, R. B., Flint, H. J., Salminen, S., Calder, P. C., & Sanders, M. E. (2014). The International Scientific Association for Probiotics and Prebiotics consensus statement on the scope and appropriate use of the term probiotic. Nature Reviews Gastroenterology & Hepatology, 11(8), 506–514. 10.1038/nrgastro.2014.66

Hu, M., Wang, D., Tang, X., Zhang, Q., Zhao, J., Mao, B., Zhang, H., & Cui, S. (2024). Improving the utilization efficiency of nitrogen source through co-culture of Lactobacillus strains with different nitrogen source metabolisms. LWT, 191, 115701. 10.1016/j.lwt.2023.115701

Jackson, R., Yao, T., Bulut, N., Cantu-Jungles, T. M., & Hamaker, B. R. (2024). Protein combined with certain dietary fibers increases butyrate production in gut microbiota fermentation. Food & Function, 15(6), 3186–3198. 10.1039/D3FO04187E

Jukic, A., Bakiri, L., Wagner, E. F., Tilg, H., & Adolph, T. E. (2021). Calprotectin: from biomarker to biological function. Gut, 70(10), 1978–1988. 10.1136/gutjnl-2021-324855

Kämpfer, A. A. M., Urbán, P., Gioria, S., Kanase, N., Stone, V., & Kinsner-Ovaskainen, A. (2017). Development of an in vitro co-culture model to mimic the human intestine in healthy and diseased state. Toxicology in Vitro, 45, 31–43. 10.1016/j.tiv.2017.08.011

Kapel, N., Ouni, H., Benahmed, N. A., & Barbot-Trystram, L. (2023). Fecal Calprotectin for the Diagnosis and Management of Inflammatory Bowel Diseases. Clinical and Translational Gastroenterology, 14(9), e00617. 10.14309/ctg.0000000000000617

Karlsson, J., Lopez-Sanchez, P., Marques, T. M., Hyötyläinen, T., Castro-Alves, V., Krona, A., & Ström, A. (2024). Physico-chemical properties of pea fibre and pea protein blends and the implications for in vitro batch fermentation using human inoculum. Food Hydrocolloids, 150, 109732. 10.1016/j.foodhyd.2024.109732

Kulich, K. R., Madisch, A., Pacini, F., Piqué, J. M., Regula, J., Van Rensburg, C. J., Újszászy, L., Carlsson, J., Halling, K., & Wiklund, I. K. (2008). Reliability and validity of the Gastrointestinal Symptom Rating Scale (GSRS) and Quality of Life in Reflux and Dyspepsia (QOLRAD) questionnaire in dyspepsia: A six-country study. Health and Quality of Life Outcomes, 6(1), 12. 10.1186/1477-7525-6-12

Levitt, D., & Levitt, M. (2018). A model of blood-ammonia homeostasis based on a quantitative analysis of nitrogen metabolism in the multiple organs involved in the production, catabolism, and excretion of ammonia in humans. *Clinical and Experimental Gastroenterology*, Volume 11, 193–215. 10.2147/CEG.S160921

Lewis, S. J., & Heaton, K. W. (1997). Stool Form Scale as a Useful Guide to Intestinal Transit Time. Scandinavian Journal of Gastroenterology, 32(9), 920–924.

Liu, M., Bayjanov, J. R., Renckens, B., Nauta, A., & Siezen, R. J. (2010). The proteolytic system of lactic acid bacteria revisited: a genomic comparison. BMC Genomics, 11(1), 36. 10.1186/1471-2164-11-36

Liu, X., Shao, J., Liao, Y.-T., Wang, L.-N., Jia, Y., Dong, P., Liu, Z., He, D., Li, C., & Zhang, X. (2023). Regulation of short-chain fatty acids in the immune system. Frontiers in Immunology, 14. 10.3389/fimmu.2023.1186892

Livsmedelsverket. (2024, December 4). Portionsguiden. https://www.livsmedelsverket.se/matvanor-halsa--miljo/matvanor---undersokningar/portionsguiden

Luzardo-Ocampo, I., & Gonzalez de Mejia, E. (2025). Plant proteins and peptides as key contributors to good health: A focus on pulses. Food Research International, 211, 116346. 10.1016/j.foodres.2025.116346

Mak, I. E. K., Yao, Y., Ng, M. T. T., & Kim, J. E. (2025). Influence of dietary protein and fiber intake interactions on the human gut microbiota composition and function: a systematic review and network meta-analysis of randomized controlled trials. Critical Reviews in Food Science and Nutrition, 1–19. 10.1080/10408398.2025.2452362

Markova, M., Koelman, L., Hornemann, S., Pivovarova, O., Sucher, S., Machann, J., Rudovich, N., Thomann, R., Schneeweiss, R., Rohn, S., Pfeiffer, A. F. H., & Aleksandrova, K. (2020). Effects of plant and animal high protein diets on immune-inflammatory biomarkers: A 6-week intervention trial. Clinical Nutrition, 39(3), 862–869. 10.1016/j.clnu.2019.03.019

Mendall, M. A., Chan, D., Patel, R., & Kumar, D. (2016). Faecal calprotectin: factors affecting levels and its potential role as a surrogate marker for risk of development of Crohn’s Disease. BMC Gastroenterology, 16(1), 126. 10.1186/s12876-016-0535-z

Mithul Aravind, S., Wichienchot, S., Tsao, R., Ramakrishnan, S., & Chakkaravarthi, S. (2021). Role of dietary polyphenols on gut microbiota, their metabolites and health benefits. Food Research International, 142, 110189. 10.1016/j.foodres.2021.110189

Moll, P., Salminen, H., Seitz, O., Schmitt, C., & Weiss, J. (2023). Characterization of soluble and insoluble fractions obtained from a commercial pea protein isolate. Journal of Dispersion Science and Technology, 44(13), 2417–2428. 10.1080/01932691.2022.2093214

Neis, E., Dejong, C., & Rensen, S. (2015). The Role of Microbial Amino Acid Metabolism in Host Metabolism. Nutrients, 7(4), 2930–2946. 10.3390/nu7042930

Parada Venegas, D., De la Fuente, M. K., Landskron, G., González, M. J., Quera, R., Dijkstra, G., Harmsen, H. J. M., Faber, K. N., & Hermoso, M. A. (2019). Short Chain Fatty Acids (SCFAs)-Mediated Gut Epithelial and Immune Regulation and Its Relevance for Inflammatory Bowel Diseases. Frontiers in Immunology, 10. 10.3389/fimmu.2019.00277

Plattner, B. J., Hong, S., Li, Y., Talavera, M. J., Dogan, H., Plattner, B. S., & Alavi, S. (2024). Use of Pea Proteins in High-Moisture Meat Analogs: Physicochemical Properties of Raw Formulations and Their Texturization Using Extrusion. Foods, 13(8), 1195. 10.3390/foods13081195

Schumacher, T., Steinmacher, T., Köster, E., Wagemans, A., Weiss, J., & Gibis, M. (2025). Physico-chemical characterization of ten commercial pea protein isolates. Food Hydrocolloids, 162, 110996. 10.1016/j.foodhyd.2024.110996

Seeburger, P., Forsman, H., Bevilacqua, G., Marques, T. M., Morales, L. O., Prado, S. B. R., Strid, Å., Hyötyläinen, T., & Castro-Alves, V. (2023). From farm to fork… and beyond! UV enhances Aryl hydrocarbon receptor-mediated activity of cruciferous vegetables in human intestinal cells upon colonic fermentation. Food Chemistry, 426, 136588. 10.1016/j.foodchem.2023.136588

Shabani, F., Mahdavi, M., Imani, M., Hosseinpour-Feizi, M., & Gheibi, N. (2020). Calprotectin (S100A8/S100A9)-induced cytotoxicity and apoptosis in human gastric cancer AGS cells: Alteration in expression levels of Bax, Bcl-2, and ERK2. Human & Experimental Toxicology, 39(8), 1031–1045. 10.1177/0960327120909530

Shah, A. B., Baiseitova, A., Zahoor, M., Ahmad, I., Ikram, M., Bakhsh, A., Shah, M. A., Ali, I., Idress, M., Ullah, R., Nasr, F. A., & Al-Zharani, M. (2024). Probiotic significance of Lactobacillus strains: a comprehensive review on health impacts, research gaps, and future prospects. Gut Microbes, 16(1). 10.1080/19490976.2024.2431643

Stamouli, S., Beber, M. E., Normark, T., Christensen, T. A., Andersson-Li, L., Borry, M., Jamy, M., & Fellows Yates, J. A. (2023). nf-core/taxprofiler: highly parallelised and flexible pipeline for metagenomic taxonomic classification and profiling. 10.1101/2023.10.20.563221

Stríz, I., & Trebichavský, I. (2004). Calprotectin - a pleiotropic molecule in acute and chronic inflammation. Physiological Research, 53(3), 245–253.

Thomas, M. S., Calle, M., & Fernandez, M. L. (2023). Healthy plant-based diets improve dyslipidemias, insulin resistance, and inflammation in metabolic syndrome. A narrative review. Advances in Nutrition, 14(1), 44–54. 10.1016/j.advnut.2022.10.002

Tiong, A. Y. J., Crawford, S., Jones, N. C., McKinley, G. H., Batchelor, W., & van ‘t Hag, L. (2024). Pea and soy protein isolate fractal gels: The role of protein composition, structure and solubility on their gelation behaviour. Food Structure, 40, 100374. 10.1016/j.foostr.2024.100374

Watzl, B. (2008). Anti-inflammatory Effects of Plant-based Foods and of their Constituents. International Journal for Vitamin and Nutrition Research, 78(6), 293–298. 10.1024/0300-9831.78.6.293

Windey, K., De Preter, V., & Verbeke, K. (2012a). Relevance of protein fermentation to gut health. Molecular Nutrition & Food Research, 56(1), 184–196. 10.1002/mnfr.201100542

Windey, K., De Preter, V., Louat, T., Schuit, F., Herman, J., Vansant, G., & Verbeke, K. (2012b). Modulation of Protein Fermentation Does Not Affect Fecal Water Toxicity: A Randomized Cross-Over Study in Healthy Subjects. PLoS ONE, 7(12), e52387. 10.1371/journal.pone.0052387

Wrong, O. M., & Vince, A. (1984). Urea and ammonia metabolism in the human large intestine. Proceedings of the Nutrition Society, 43(1), 77–86. 10.1079/PNS19840030

Wu, S., Bhat, Z., Gounder, R., Mohamed Ahmed, I., Al-Juhaimi, F., Ding, Y., & Bekhit, A. (2022). Effect of Dietary Protein and Processing on Gut Microbiota—A Systematic Review. Nutrients, 14(3), 453. 10.3390/nu14030453

Yao, Y., Cai, X., Fei, W., Ye, Y., Zhao, M., & Zheng, C. (2022). The role of short-chain fatty acids in immunity, inflammation and metabolism. Critical Reviews in Food Science and Nutrition, 62(1), 1–12. 10.1080/10408398.2020.1854675

Zhou, M., Ma, J., Kang, M., Tang, W., Xia, S., Yin, J., & Yin, Y. (2024). Flavonoids, gut microbiota, and host lipid metabolism. Engineering in Life Sciences, 24(5), 2300065. 10.1002/elsc.202300065

Zhu, H.-G., Tang, H.-Q., Cheng, Y.-Q., Li, Z.-G., & Tong, L.-T. (2021). Potential of preparing meat analogue by functional dry and wet pea (Pisum sativum) protein isolate. LWT, 148, 111702. 10.1016/j.lwt.2021.111702

