## Supplementary file for "Effects of incrementally increased plant-based protein intake on gut microbiota and inflammatory–metabolic biomarkers in healthy adults"

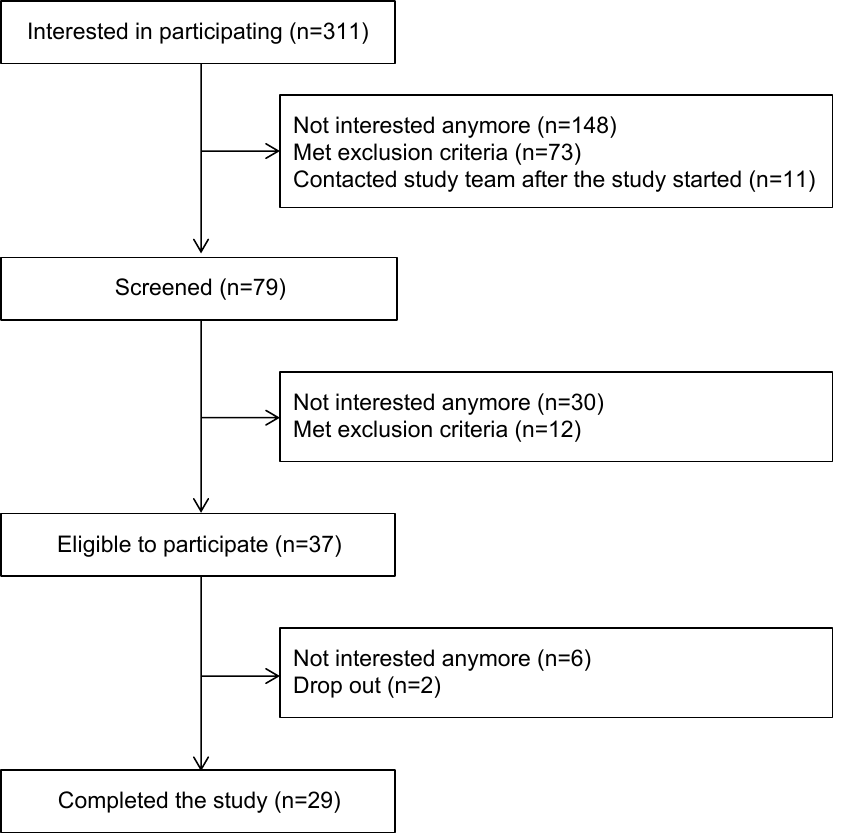

**Supplementary Figure 1. Study Workflow.**

**Supplementary Table 1. Chemical Composition of the Provided Isolated Pea Protein.**

|  | **Week 1** | **Week 2** | **Week 3** | **Week 4** |
| --- | --- | --- | --- | --- |
|  | ***g*** | ***g*** | ***g*** | ***g*** |
| **Isolated Pea Components** |  |  |  |  |
| Protein* | 17.325 | 34.650 | 51.975 | 69.300 |
| Starch | 0.041 | 0.081 | 0.122 | 0.162 |
| Fat | 1.156 | 2.313 | 3.469 | 4.625 |
| Fibre NDF | 0.162 | 0.325 | 0.487 | 0.649 |
| Fibre ADF | 0.041 | 0.081 | 0.122 | 0.162 |
| **Essencial Amino Acids** |  |  |  |  |
| Histidine | 0.366 | 0.733 | 1.099 | 1.466 |
| Isoleucine | 0.573 | 1.147 | 1.720 | 2.294 |
| Leucine | 1.322 | 2.644 | 3.967 | 5.289 |
| Lysine | 1.274 | 2.549 | 3.823 | 5.098 |
| Phenylalanine | 0.860 | 1.720 | 2.581 | 3.441 |
| Threonine | 0.637 | 1.274 | 1.912 | 2.549 |
| Valine | 0.637 | 1.274 | 1.912 | 2.549 |
| **Non-essencial Amino Acids** |  |  |  |  |
| Alanine | 0.685 | 1.370 | 2.055 | 2.740 |
| Arginine | 1.131 | 2.262 | 3.393 | 4.524 |
| Aspartic acid | 2.151 | 4.301 | 6.452 | 8.602 |
| Cysteine | 0.048 | 0.096 | 0.143 | 0.191 |
| Glutamic acid | 3.106 | 6.213 | 9.319 | 12.426 |
| Glycine | 0.637 | 1.274 | 1.912 | 2.549 |
| Proline | 0.797 | 1.593 | 2.390 | 3.186 |
| Serine | 0.956 | 1.912 | 2.867 | 3.823 |
| Tyrosine | 0.605 | 1.211 | 1.816 | 2.421 |

*Based on the median bodyweight of 69.3 kg. Results are calculated based on Auer et al. 2024. Neutral Detergent Fiber (NDF) and Acid Detergent Fiber (ADF).

**Supplementary Table 2. Gastrointestinal Symptom Rating Scale Scores. Bristol Stool Scale Scores and Physical Activity.**

|  | **Week 4** |  | **Week 5** |  | **Week 6** |  | **Week 7** |  | **Week 8** |  |
| --- | --- | --- | --- | --- | --- | --- | --- | --- | --- | --- |
|  | *Median (IQR)* | *n* | *Median (IQR)* | *n* | *Median (IQR)* | *n* | *Median (IQR)* | *n* | *Median (IQR)* | *n* |
| **GSRS Score** |  |  |  |  |  |  |  |  |  |  |
| Diarrhoea | 1.0 (1.0-1.5) | 29 | 1.0 (1.0-2.0) | 29 | 1.0 (1.0-1.7) | 29 | 1.3 (1.0-1.7) | 29 | 1.0 (1.0-1.7) | 29 |
| Indigestion | 1.8 (1.1-2.3) | 29 | 1.8 (1.1-2.4) | 29 | 1.5 (1.1-2.5) | 29 | 1.8 (1.1-2.3) | 29 | 2.0 (1.1-2.7) | 29 |
| Constipation | 1.0 (1.00-1.7) | 29 | 1.0 (1.0-1.3) | 29 | 1.0 (1.0-1.3) | 29 | 1.3 (1.0-1.9) | 29 | 1.3 (1.0-1.7) | 29 |
| Abdominal pain | 1.3 (1.0-1.7) | 29 | 1.0 (1.0-1.7) | 29 | 1.0 (1.0-1.5) | 29 | 1.0 (1.0-1.7) | 29 | 1.3 (1.0-1.9) | 29 |
| Reflux | 1.0 (1.0-1.1) | 29 | 1.0 (1.0-1.2) | 29 | 1.0 (1.0-1.2) | 29 | 1.0 (1.0-1.1) | 29 | 1.0 (1.0-1.1) | 29 |
| Total score | 1.3 (1.0-1.7) | 29 | 1.4 (1.1-1.7) | 29 | 1.3 (1.1-1.6) | 29 | 1.5 (1.1-1.8) | 29 | 1.5 (1.0-2.0) | 29 |
| **BSS Score** |  |  |  |  |  |  |  |  |  |  |
| Type/wk | 3.8 (3.4-4.3) | 28 | 4.3 (3.4-5.0) | 28 | 3.8 (3.4-5.0) | 28 | 4.0 (3.4-4.8) | 28 | 3.9 (3.2-4.7) | 28 |
| Frequency/d | 0.9 (0.7-1.3) | 29 | 1.1 (0.9-1.3) | 29 | 1.0 (0.8-1.2) | 29 | 1.0 (0.8-1.3) | 29 | 1.0 (0.7-1.3) | 29 |
| **PA** |  |  |  |  |  |  |  |  |  |  |
| MET min/wk^**^ | 2461 (1655-3920) | 28 | 2432 (1598-4372) | 28 | 2245 (1455-3390) | 28 | 2062 (1003-3167) | 28 | 1951 (1134-3461) | 28 |
| PAL/wk | 2.0 (2.0-3.0) | 28 | 3.0 (2.0-3.0) | 28 | 3.0 (2.0-3.0) | 28 | 2.4 (2.0-3.0) | 28 | 2.0 (2.0-3.0) | 28 |

Data is shown as median (IQR) and includes results from a Friedman test with Dunn’s multiple comparisons of each intervention week vs. the baseline (week 4); ^**^p<0.01; GSRS: Gastrointestinal Symptom Rating Scale. BSS: Bristol Stool Scale. PA: Physical Activity. PAL: Physical Activity Level.
